# A Cre-amplifier to generate and detect genetic mosaics *in vivo*

**DOI:** 10.1101/715490

**Authors:** Francesco Trovato, Riccardo Parra, Enrico Pracucci, Silvia Landi, Olga Cozzolino, Gabriele Nardi, Federica Cruciani, Laura Mosti, Andrzej Cwetsch, Laura Cancedda, Laura Gritti, Carlo Sala, Chiara Verpelli, Andrea Maset, Claudia Lodovichi, Gian Michele Ratto

**Affiliations:** National Enterprise for Nanoscience and Nanotechnology (NEST), Istituto Nanoscienze Consiglio Nazionale delle Ricerche (CNR) and Scuola Normale Superiore Pisa, 56127 Pisa, Italy; Institute of Neuroscience CNR, Pisa, Italy; Istituto Italiano di Tecnologia, Genoa, Italy; Università degli studi di Genova, Genoa, Italy; Istituto Telethon Dulbecco; Institute of Neuroscience CNR, Milan, Italy; Veneto Institute of Molecular Medicine, Padua, Italy; Institute of Neuroscience CNR, Padua, Italy

**Author notes:** **Author’s contribution.** The study was designed by FT, RP, GMR and was supervised by FT and GMR. Beatrix was designed and generated by FT and RP with the help of LM. Early versions of Beatrix have been extensively tested *in vitro* and *in vivo* by LC, AC, CL, CS, CV and LG. *In utero* electroporations of wild type animals were performed by LC, AC, FT, OC, SL, EP, and GN. Experiments on PTEN strain were performed by FT, EP, SL, OC, GN, FC. Experiments on cultures of MeCP2-KO mice were performed by CS, CV and LG. Experiments on the olfactory bulb of PTEN mice were performed by CL and AM and FC helped with data acquisition. The paper was written by FT, GMR, EP, OC and GN with furthers contributions from all authors. Please send all correspondence to Francesco Trovato and Gian Michele Ratto.

## Abstract

Cre-Lox manipulation is the gold standard for cell-specific expression or knockout of selected genes. However, it is not unusual to deal with conditions of low Cre expression or transient activation, which can often go undetected by conventional Cre-reporters. We designed Beatrix, a general-purpose tool specifically devised to amplify weak Cre recombinase activity, and we used it to develop a powerful approach for the *in vivo* generation and detection of sparse mosaics of mutant and wild type cells.

## Main

Genetic mosaicism refers to the presence of genetically distinct cellular populations within the same individual. Mosaic modelling is a major theme in life science, as mosaicism is associated to many pathological conditions: several Mendelian disorders, chromosomal aberrations, neurological dysfunctions as focal dysplasia or autism spectrum disorders, and cancer have been directly or indirectly connected to certain degrees of mosaicism^1–3^. Moreover, sparse mosaic labelling is a powerful tool for the study of single-cell functions and cell-autonomous effects of selective overexpression/knockout. Currently, many diffused approaches for the generation of mosaic models make use of the Cre-Lox technology. In these systems, Cre-activated reporters are used to tell recombinant and non-recombinant cells apart^4–7^.

Our goal is to define a Cre-based strategy to differentially label both wild type (WT) and knockout cells (KO) for a gene of interest, with a fine control of the mosaicism level. The general idea is to co-transfect two different plasmids. The first plasmid, based on the Cre-switch design^8^, is a Cre reporter relying on the FLEx system^9^ that switches expression between two fluorophores (RFP/GFP) depending on Cre activity. The second plasmid, carries Cre recombinase and it acts as a trigger for the recombination. The concentration of the trigger plasmid determines the mosaicism entity. However, the necessity to have a biunivocal correspondence between the genomic floxed allele status and the Cre-reporter readout is difficult to fulfill with the low levels of recombinase activity required to induce a sparse recombination. Indeed, at low Cre concentration, recombination has been described to occur in the reporter, but not in the gene of interest or *vice versa*, leading to false positive or false negative cells respectively^10,11^. Thus, the incomplete recombination in cells experiencing a low Cre titer results in uncertainty about the floxed allele status, witnessed by the coexistence of both the red and the green forms of the Cre-switch plasmid copies within cells (Fig. 1a).

**Fig. 1.**
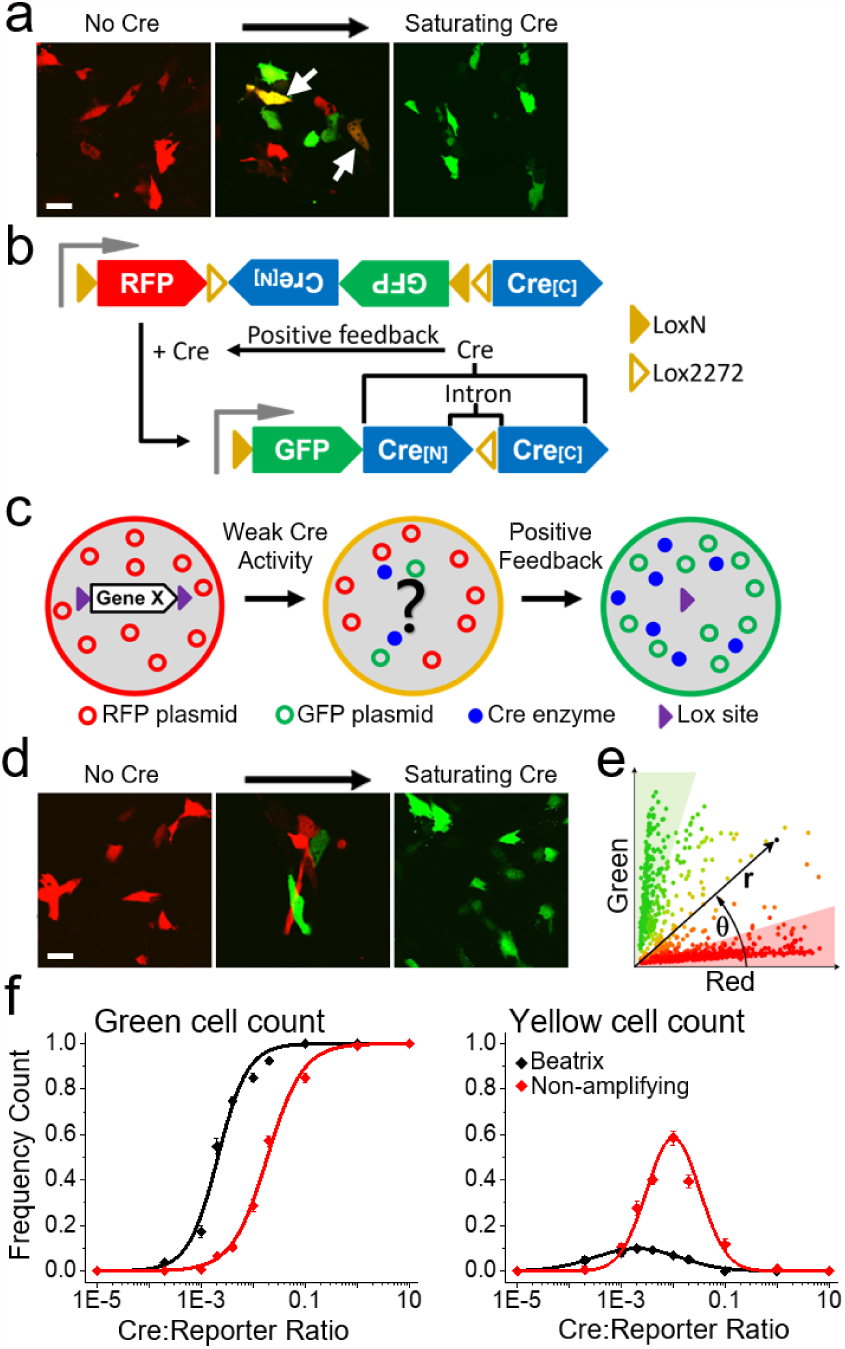
Beatrix: a sensor and effector of Cre activity. **a**, Representative two-photon images of NIH3T3 cells transfected with a non-amplifying Cre-switch based double reporter at three different concentrations of the Cre trigger. At low trigger plasmid titer (central panel) the mosaic includes a significant number of yellow cells (white arrows). Scale bar 20 µm. **b**, Design of the fluorescent sensor and of the Cre amplifier embedded in Beatrix. See Supplementary Fig.1 for the recombination sequence. **c**, Schematic representation of the operative mechanism of the sensor. **d**, NIH3T3 cells transfected with Beatrix at three different molar ratios of the Cre trigger. The central panel shows the effect of the amplification at low concentration of the Cre trigger. Scale bar 20 µm. **e**, Polar coordinates representation of the red and green fluorescence. The radius r measures the overall fluorescence intensity. The polar angle θ measures the hue of each cell, proportional to the ratio between red and green fluorescence (see Methods, Supplementary Data). Cells lying in green and shaded areas are expressing either the fully recombined plasmid (green) or the native plasmid (red). All other cells are considered partially recombined (yellow). **f**, The left panel shows the relative frequency of cells that express only GFP as a function of the Cre:Reporter plasmid ratio for Beatrix (black) and the non-amplifying control (red). The right panel shows the relative frequency of cells presenting mixed expression of GFP and RFP.

To overcome this major limitation we developed Beatrix, an amplifier of Cre activity sensitive to low Cre concentration, which embeds a Cre-dependent Cre gene copy within a FLEx structure (Fig.1 b). In absence of Cre activity, Beatrix drives the exclusive expression of the RFP protein (dsRed2). Upon Cre-mediated recombination, the Beatrix cassette rearrangement leads to the RFP excision, and the expression of both GFP and Cre (Fig. 1b and Supplementary Fig. 1 and 2).

Like in a domino chain reaction, this method exploits the high plasmid copy number of Beatrix into transfected cells and its inherent positive feedback to amplify a weak Cre trigger into a saturating Cre activity (Fig. 1c). Accordingly, RFP signal will correspond to completely absent Cre, while GFP signal to a strong Cre expression. We initially explored simpler variants of our construct where the whole intron-interrupted Cre gene was inserted in antisense in the Cre-switch architecture. Unfortunately, all of these constructs presented a substantial leakage due to self-activation of the embedded Cre amplifier^12^, and we eventually determined that the key breakthrough was the division of the Cre gene in two exons assembled in opposite orientations (Supplementary Fig. 3).

Fig. 1d shows how, in presence of the amplification provided by Beatrix, cells appear clearly segregated in red and green cohorts. We quantified the difference between a conventional Cre-switch reporter and Beatrix by looking at the number of green and yellow cells over the total transfected population as a function of the Cre trigger titer, adopting the criteria depicted in Fig. 1e. Fig. 1f shows the frequency of cells expressing only GFP in different mosaics. The embedded amplification strongly enhances the sensitivity of Beatrix to low trigger plasmid titers, shifting the curve toward a tenfold lower concentration without affecting the slope. At the same time, the frequency of recombinant cells with uncertain status (yellow cells) is massively reduced. Prompted by this result, we tested our approach for the generation of mosaics *in vivo* in the mouse brain after *in utero* electroporation^13^ (IUE) with Beatrix and variable concentrations of the Cre plasmid (Supplementary Fig. 4). Fig. 2a shows representative fields of neuronal mosaics in layer 2/3 (L2/3) of mouse cortex acquired *in vivo* at P30 after IUE with Cre and either Beatrix or the non-amplifying reporter at 1:150 molar ratio. We assessed these mosaics by quantifying the fluorescence signal of each neuron (Fig 2b).

**Fig. 2.**
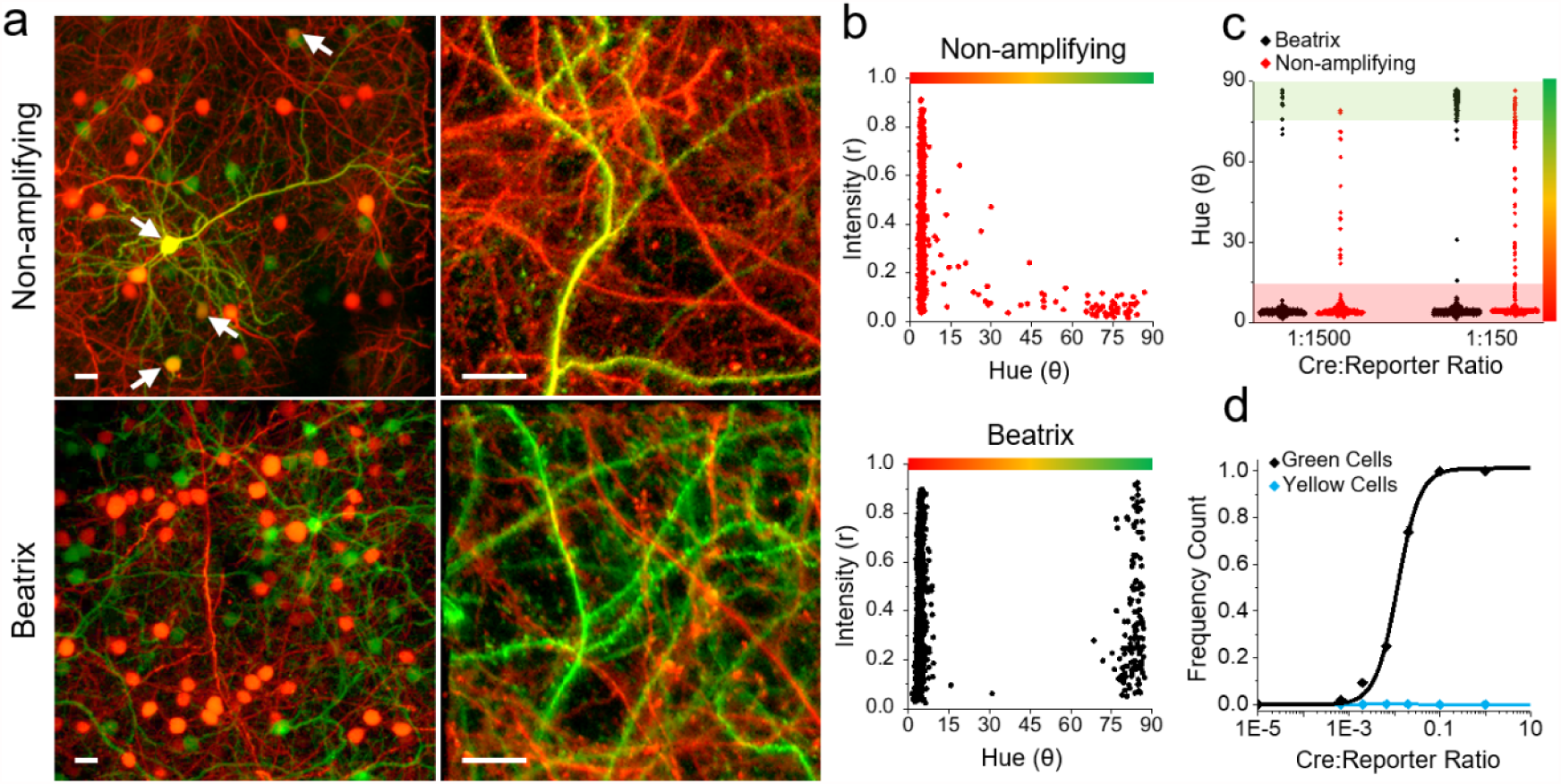
Comparison of *in vivo* expression mosaic induced with Beatrix or with a conventional reporter. **a**, Left panels show representative fields of expression mosaic obtained with a Cre:Reporter ratio of 1:150. The arrowheads indicate cell bodies expressing both RFP and GFP that are frequent in the mosaic obtained without Cre amplification. The information about the recombination state of the neuron can also be obtained by examining the dendrites (right panels). In absence of amplification, dendrites show a uniform co-localization of GFP and RFP, which is absent in presence of amplification (see Supplementary Fig. S4). **b**, Cumulative data obtained from 4 mice transfected with Beatrix and Cre at the 1:150 ratio (864 neurons, black symbols on top) and from 3 mice transfected with a non-amplifying reporter and Cre (609 cells, red symbols on bottom); these plots show the effect of the amplification on binarization (polar angle θ) and mean fluorescence intensity (radial distance r). In absence of amplification the strong negative correlation between intensity and hue is confirmed by Pearson correlation coefficient R=-0.45 (F-test p<10^−4^). **c**, Scatterplot showing the difference in hue variation upon a tenfold dilution of the Cre controller using Beatrix or the non-amplifying reporter. Cells show a very strong binarization only in presence of amplification. 1:1500 ratio (n=3 mice per group), 1:150 ratio (n=4, Beatrix; n=3 no amplification). **d**, Frequency count of cells expressing only GFP (black) or both RFP and GFP (cyan) as a function of the Cre:Beatrix plasmid ratio. The number of yellow cells is negligible for every Cre:Beatrix plasmid ratio.

In absence of amplification, mosaics are affected by a considerable number of partially recombined yellow cell bodies and dendrites, independently of the mosaicism level (Fig 2b,c).

Conversely, the mosaics induced by Beatrix are virtually perfectly binarized, with a negligible presence of partially recombined cells (less than 0.1%). Importantly, the segregation of neurons in the two classes is independent on the concentration of the triggering Cre plasmid. Thus, changes in concentration of the Cre trigger plasmid result only in a tuning of green to red cells number, without any significant increase in the yellow cells component (Fig. 2d). A further notable feature of the mosaics induced by Beatrix is the high mean fluorescence intensity of the transfected cells, regardless of their recombination status. By contrast, our data show how the non-amplifying reporter presents a clear correlation between fluorescence intensity and hue, as cell fluorescence intensity progressively decreases going from red to green (Fig. 2b).

As a proof-of-principle demonstration of our method, we generated a sparse mosaic of expression of the PTEN gene in a conditional knockout mouse strain (Pten^flox^). PTEN (Phosphatase and tensin homolog) is a phosphatase that negatively regulates the PI3K/AKT/mTOR pathway exerting an important role in the regulation of several crucial cellular processes. The loss of PTEN plays a pivotal role in cancer progression, non-cancerous neoplasia, focal cortical dysplasia and has been associated with autism spectrum disorders, therefore providing multiple points of interest for mosaicism modelling^14–17^. In this set of experiments we used Beatrix in two different contexts: in cortical pyramidal neurons after IUE and in the granule cell neurons of the olfactory bulb (OB) after post-natal electroporation (Fig 3a, Supplementary Fig. 5 and 6). We first focused on the validation of Beatrix as a reporter of the recombination status of the target gene, by immunofluorescence against the PTEN protein. Quantitative colocalization analysis between the immunostaining and the reporters confirms that the recombination of Beatrix is always associated to the loss of the target gene staining. Symmetrically, all red cells are positive to the PTEN immunostaining proving the absence of false positives and false negatives in the mosaic (Fig. 3b, Supplementary Fig. 6). Next, we studied the consequences of PTEN KO at the single cell level. GFP^+^ neurons exhibit hypertrophic soma and dendrites with respect to the RFP^+^ internal controls (Fig. 3c, Supplementary Fig. 5 and 6). Furthermore, the PTEN-defective pyramidal neurons presented an aberrant migratory phenotype, as shown in Fig. 3d and Supplementary Fig. 5d. Interestingly, the KO cells cohort is characterized by a deeper localization of cell bodies and a larger dispersion of their depth, which is consistent to previous observations^18^, and with clinical manifestation of focal cortical dysplasia in humans^19^. Importantly, this effect seems to be in large measure cell-autonomous, since the RFP^+^ neurons show a normal layering. Fig. 3e and Supplementary Fig. 5b,c show how the KO neurons are characterized by a strong increase in spine density, with a marked difference in primary and secondary branches and in apical dendrites, with respect to the WT controls. Moreover, dendrites present marked morphological abnormalities in PTEN-deficient neurons, with a generally higher presence of immature spines (Supplementary Fig. 5 and 6). The immature phenotype of dendrites is also highlighted by the high degree of residual structural plasticity observed by *in vivo* time lapse imaging at P25 (Supplementary Fig. 5e), an age at which the somatosensory cortex should be almost completely stabilized^20^. Notably, we also report the occasional occurrence of somatic spines in KO cells, which are not present in any of the WT somata in the acquired fields (Supplementary Fig. 5b). Interestingly inhibitory granule cells of the OB present a different pattern of spine distribution, with the highest effect on spine density in the basal and proximal compartments (Supplementary Fig. 6). Finally, to further validate our approach in a different experimental paradigm, we also induced a KO mosaic of the MeCP2 gene on cultured cortical neurons (Supplementary Fig. 7).

**Fig. 3.**
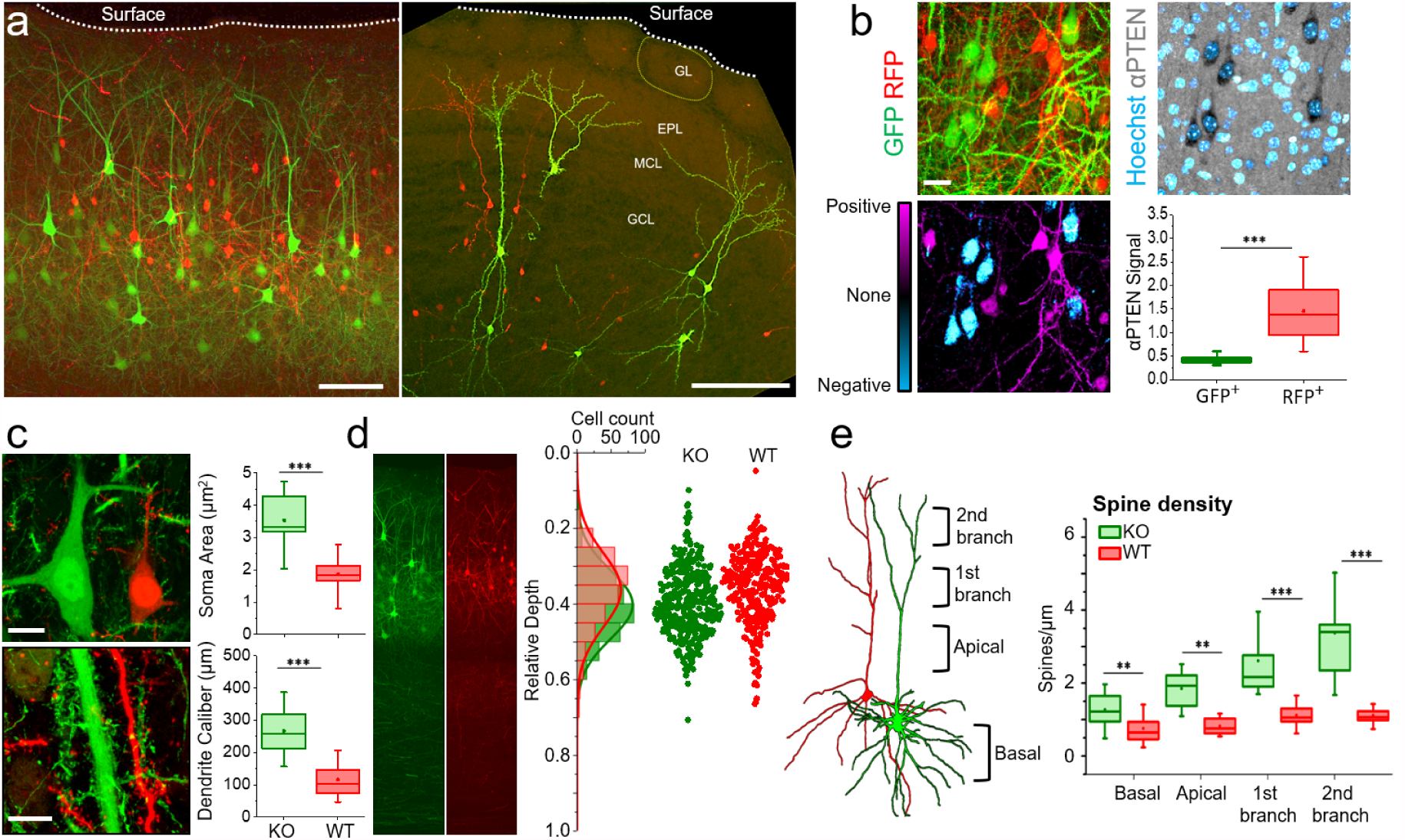
Beatrix-induced mosaic in a Pten^Flox^ conditional mouse. **a**, Maximum intensity projection of a Pten^Flox^ KO mosaic in mouse cortex and olfactory bulb (GL=glomerular layer; EPL=external plexiform layer; MCL= mitral cell layer; GCL= granule cell layer). Scale bar 100 µm. **b**, The lack of PTEN was confirmed by performing immunostaining against PTEN in the cortex mosaic (upper panels). The colocalization analysis between Beatrix reporters and PTEN immunofluorescence highlights the absolute reliability of the system as a reporter of the genomic status (bottom left). This is confirmed by the quantification of immunostaining signal in RFP/GFP positive cells (bottom right; n=3 mice). **c**, Structural details from PTEN knockout mosaics showing that the loss of PTEN causes hypertrophy of both somata and dendrites of cortical pyramidal neurons. Scale bar 10 µm. **d**, The distribution of the relative depth of neurons shows that the embryonic deletion of PTEN causes a defective migration following neurogenesis, since KO cells show a larger dispersion between the cortical layers. **e**, Spine density was quantified in different cellular compartments, highlighting a strong increase in spinogenesis, especially in the apical compartment (n=3 mice).

Although other methods for mosaic analysis^4–7^ and increased reporting reliability^5,12^ have previously been proposed, the method we present here overcomes the trade-off among sparseness, brightness and reporting reliability in an all-in-one tool, without the need to combine multiple biotechnological systems. Moreover, the use of Beatrix is not limited to the generation of sparse mosaics. Its potential includes every situation where weak or transient Cre activation represents a source of failure of detection. An important case in point involves the use of the tamoxifen-inducible form of Cre (Supplementary Fig. 8 and 9). Taken together, these results confirm the robustness of our tool and its extensibility to various conditions, proving Beatrix as a novel experimental asset to understand the physio-pathological relevance of mosaicism at the single cell level and to extend the reporting reliability in currently available Cre-based systems, animal strains or vectors.

## Supplementary Data

**Supplementary Fig. 1.**
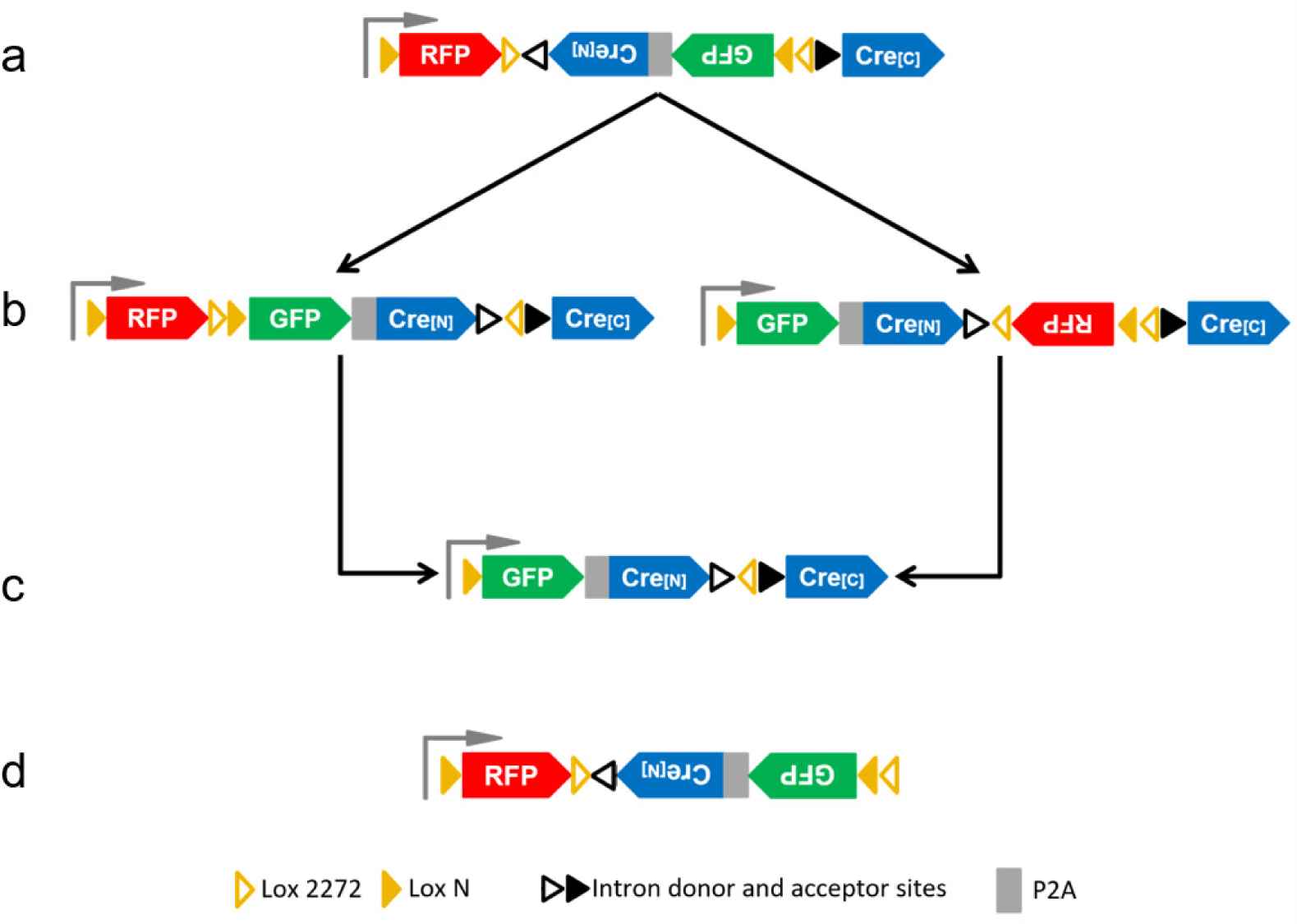
Structure and recombination of Beatrix in the presence of Cre. **a**, Schematic structure of Beatrix. Two steps of Cre-driven recombination are required to transform the RFP-expressing Beatrix, to the fully recombined GFP and Cre-expressing plasmid. **b**, The first step of recombination produces two alternative intermediate structures, depending on which lox pair is recombined first. The recombination of the Lox 2272 sites produces the intermediate shown on the left while the recombination of the Lox N sites leads to the structure shown on the right. **c**, The second recombination leads to the excision of RFP and the production of a structure that expresses only GFP and Cre. Notably, after the full recombination of Beatrix, only one Lox 2272 and one Lox N site are present, making any other recombination extremely unlikely. **d**, Structure of the control reporter: this plasmid includes the same lox sites as Beatrix, but the exon 2 of Cre gene has been removed. Therefore, it cannot lead to the amplification of Cre activity.

**Supplementary Fig. 2.**
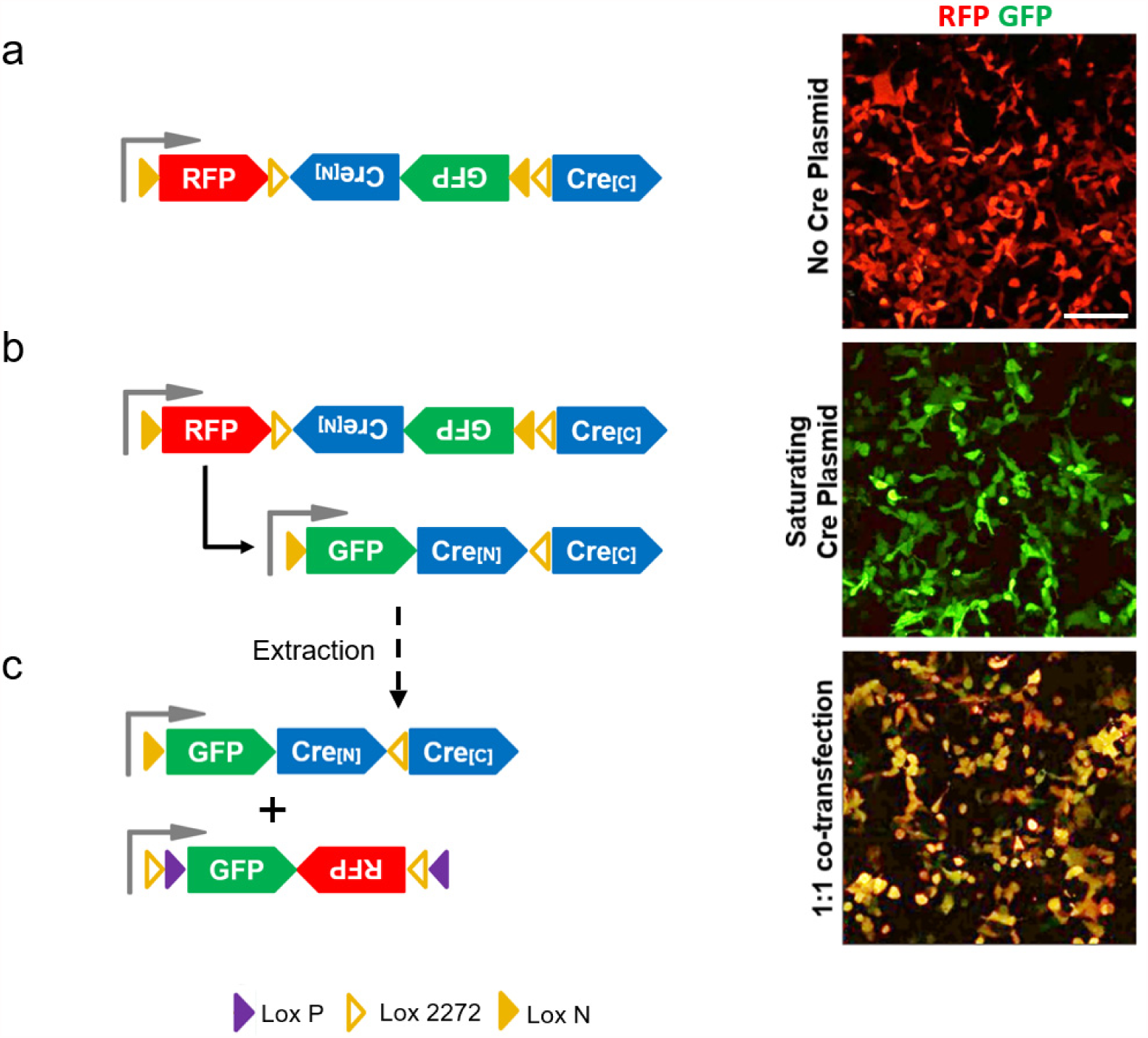
The product of Beatrix recombination is a fully functional Cre recombinase. **a**, The transfection of HEK 293T cells with Beatrix without the Cre controller produces a uniform population of cells expressing RFP only. **b**, Cells were co-transfected with Beatrix and a saturating concentration of Cre-expressing plasmid. This produced a full recombination of Beatrix thus leading to the expression of GFP only and of the reconstituted Cre. The DNA was extracted from this culture and we isolated and sequenced the fully recombined Beatrix thus confirming the expected structure depicted by the lower scheme. **c**, To prove the efficacy of Cre-activity supported by the recombined Beatrix, we used a simple Cre-switch reporter that expresses GFP in absence of Cre activity and, in presence of Cre, undergoes a two-step recombination involving Lox 2272 (yellow hollow triangles in the lower scheme) and Lox P sites (purple triangles) leading to the production of RFP. Purified Beatrix extracted from the culture shown in **b**, was co-transfected with the reporter in a 1:1 ratio. After transfection, all cells expresses both fluorescent proteins: the GFP provided by Beatrix and the RFP that is the reporter of the Cre-mediated recombination. Scale bar 50 µm.

**Supplementary Fig. 3.**
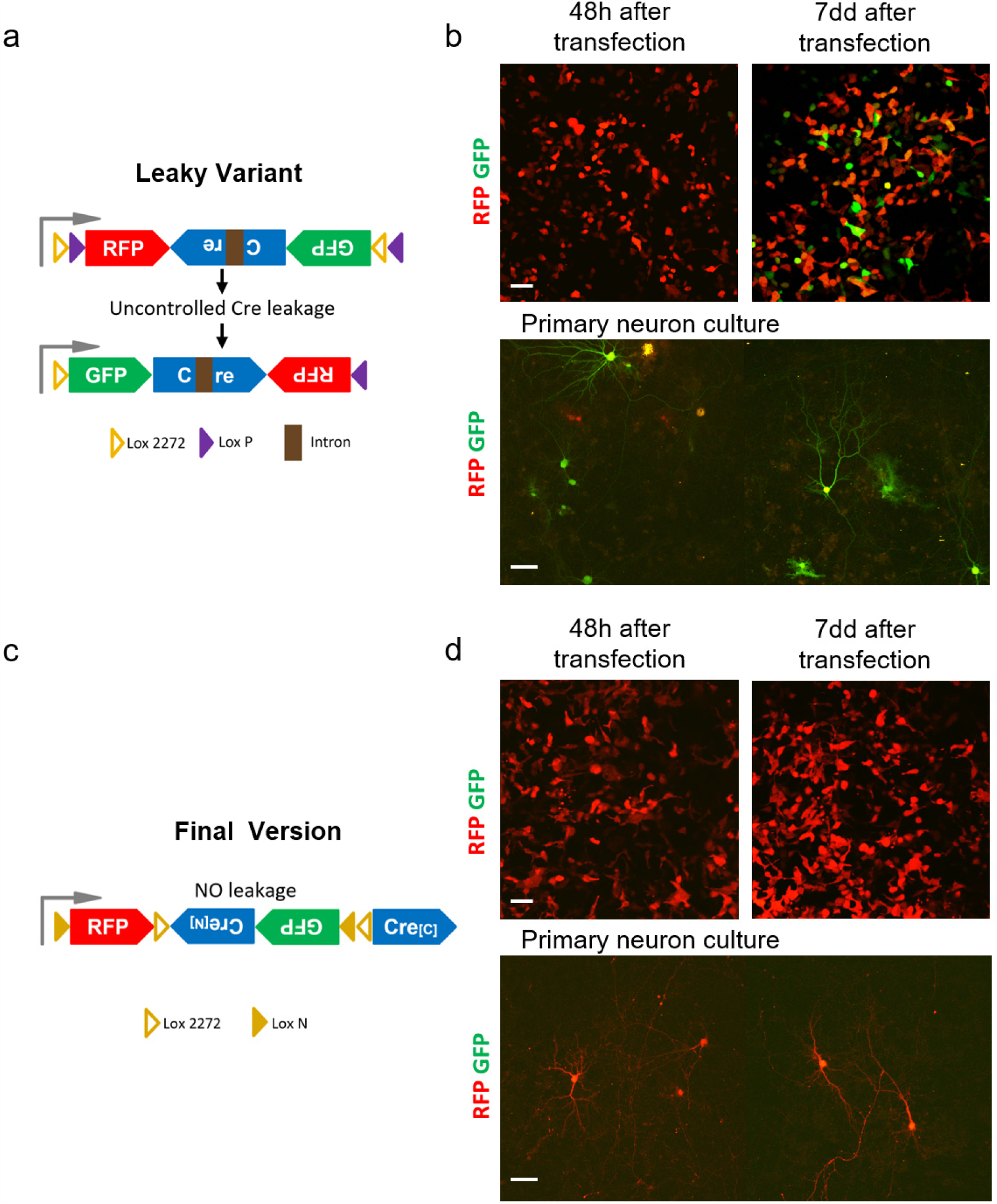
A previous variant of Beatrix showed a high non-induced self-activation level. **a**, An early version of the Beatrix construct was designed by inserting a GFP-P2A-Cre unit in antisense orientation using as a parent structure a Cre-Switch reporter (Addgene #37120). This plasmid should express an RFP before recombination and both GFP and Cre upon recombination. Unfortunately, this architecture suffers of an uncontrollable self-activation of the plasmid in absence of any Cre induction most likely due to antisense transcription of the Cre gene. Thus, a proper use of this construct is impossible since it is affected by uncontrollable leakage. This effect has also been reported in an analogous structure described recently (Fernández-Chacón, M. et al. 2019). **b**, 48 hours after transfection in HEK293T cells this construct resulted almost exclusively in RFP expressing cells (upper left panel) but, after 7 days from transfection, the leakage was very prominent and the amplification process was started in a large number of cells that switched from red to green (upper right panel). The leakage of this construct is more prominent in primary cultures of cortical neurons (bottom panel). **c**, The final Beatrix structure is characterized by the separation of the Cre gene in two exons that are orientated in opposite directions, preventing any unwanted transcription of the recombinase before induction. **d**, By adopting this strategy leakage dropped to zero and when transfected in both HEK293T and primary cortical neurons the sensor resulted exclusively in RFP expression. Scale bar 50 μm.

**Supplementary Fig. 4.**
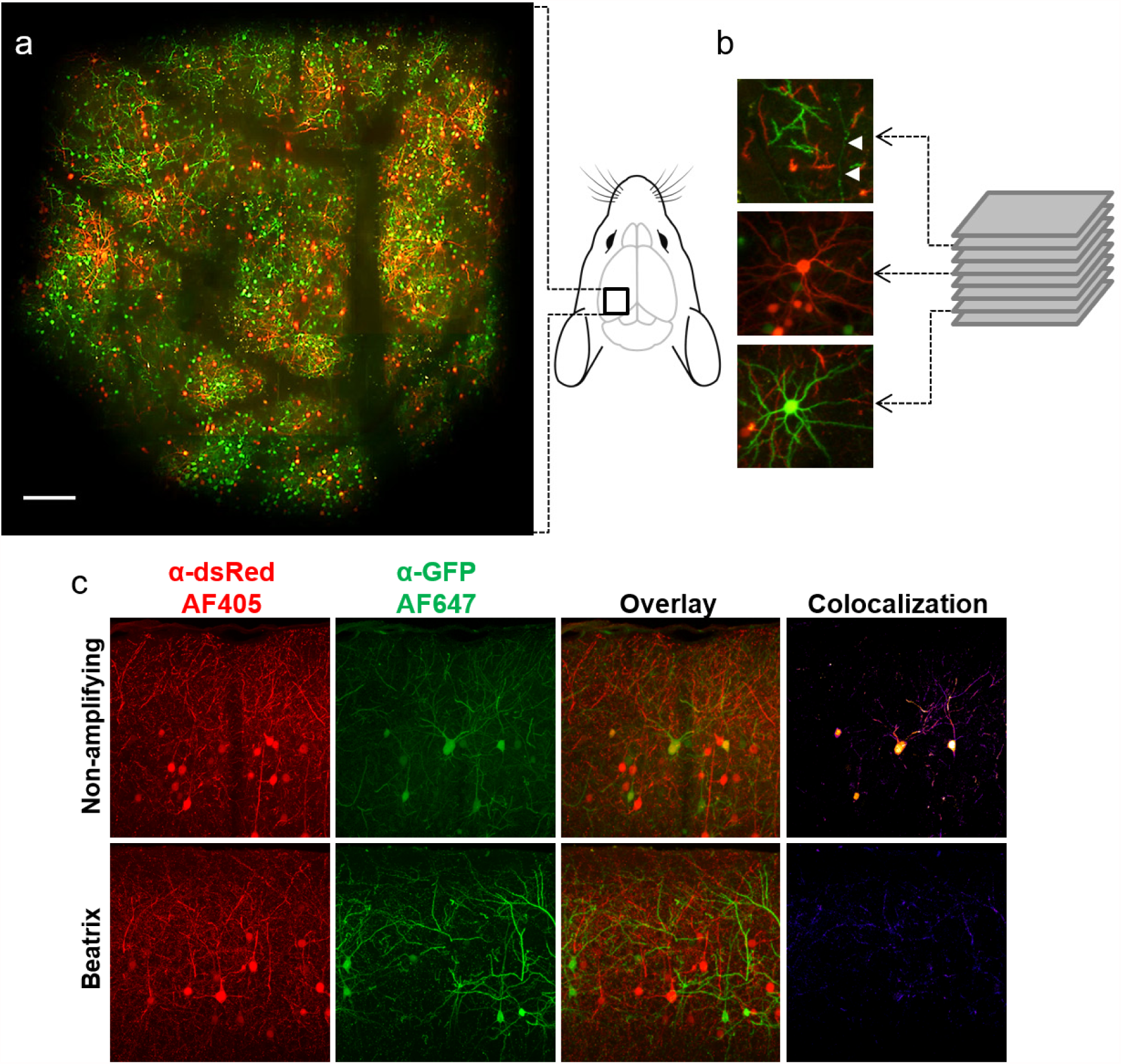
Two photon *in vivo* imaging of Beatrix-induced mosaics obtained by *in utero*electroporation. **a**, Wide field mosaic showing the entire area enclosed by the cranial window. The image obtained at P30 is a composite maximum projection of a stack imaged every 10 μm from the surface down to about 250 µm depth. All transfected cells are pyramidal neurons belonging to layers 2,3. Scale bar 200 µm. **b**, Details relative to fields placed at 25, 130 and 250 µm depth (in clockwise direction). The white arrows point to a superficial axon in layer 1 running through a field of apical dendrites. Fields are 100 µm wide. **c**, Anti-RFP and anti-GFP immunostaining of cortical slices obtained by mosaic models. The upper images show the immunofluorescence of α-dsRed and α-GFP in a mosaic generated with the non-amplifying reporter. The two proteins show a high colocalization degree indicating the partial recombination of the plasmids. The lower panels show the complete lack of colocalization in mosaics generated with Beatrix. Scale bar 50 μm.

**Supplementary Fig. 5.**
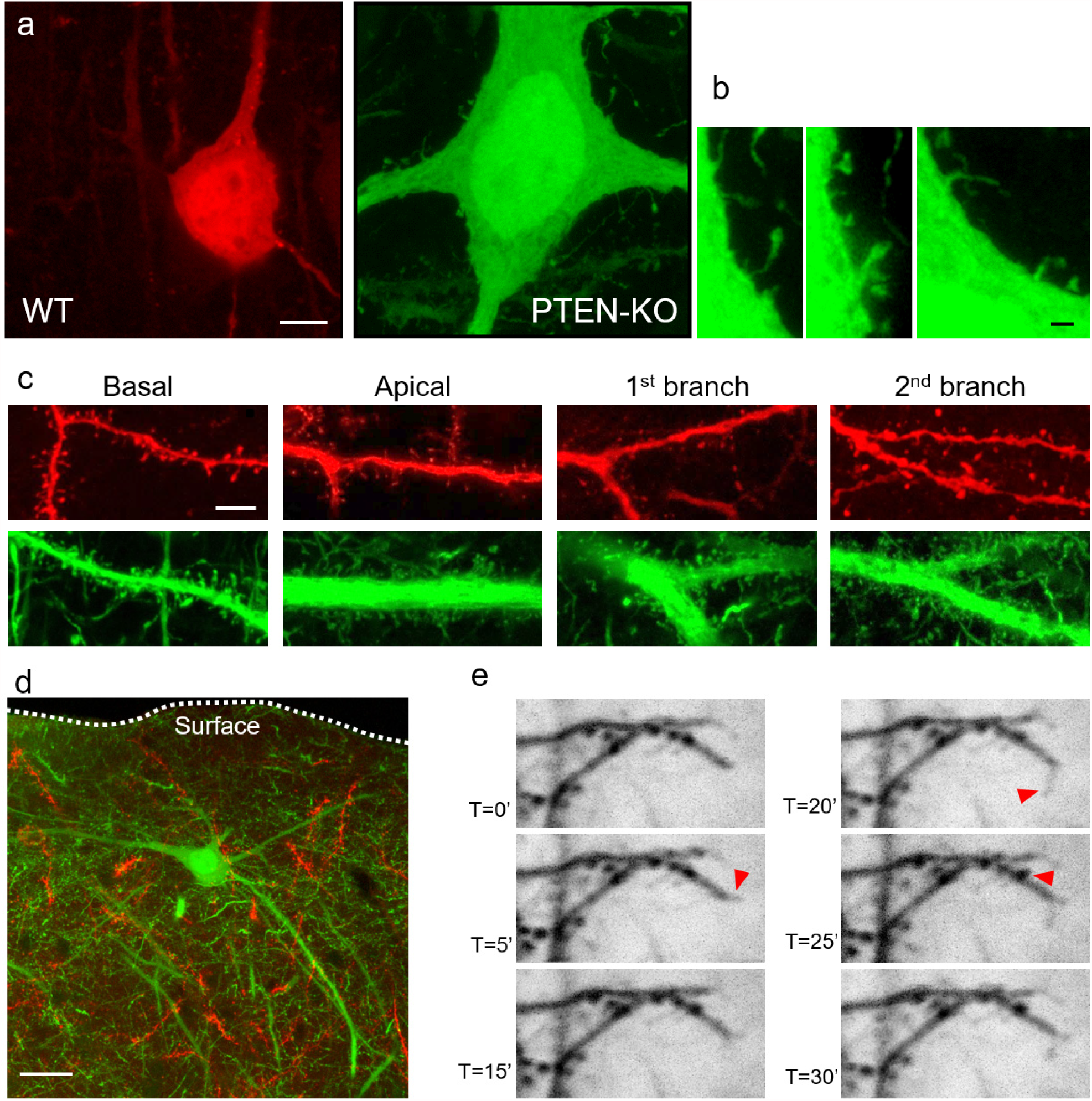
Details from the PTEN-KO mosaic model. Two photon *in vivo* imaging of Beatrix-induced mosaics obtained by *in utero* electroporation and imaged at P30 in the somatosensory cortex. **a**, Cell bodies of a wild type and of a hypertrophic KO pyramidal neuron. Scale bar 5 µm. **b**, High magnification images show the ectopic presence of dendritic spines and filopodia on the cell body surface. Scale bar 1 µm. **c**, Representative images of dendritic compartments from wild type (upper panels) and PTEN knockout pyramidal neurons (lower panels). The images show how KO neurons exhibit the hypertrophic phenotype in all cellular compartments, along with a substantial increase in spine density and filopodia numbers (as quantified in Fig. 3e). Scalebar 5 µm **d**, The alteration of the migratory phenotype induced by PTEN knockout in neuronal precursors occasionally results in a complete displacement of neurons. This image shows a cortical pyramidal neuron located in layer 1, approximately 40 µm from the brain surface, projecting its dendrites perpendicularly to the correct orientation. Scalebar 20 µm. **e**, *In vivo* time lapse imaging of dendritic spines and of two dendrite terminals in layer 2 of the somatosensory cortex. These dendrites, shown in black for clarity, are PTEN-KO since they express GFP only. Imaging was performed at P25 and at this age it is expected that dendrites have reached a mature structure characterized by very limited short time spine motility and dendritic growth. These data show a tantalizing amount of plasticity, possibly associated to immature phenotype of the PTEN-KO dendrites also witnessed by the large number of filopodia as shown in panel c. Arrowheads point to structures of high motility.

**Supplementary Fig. 6.**
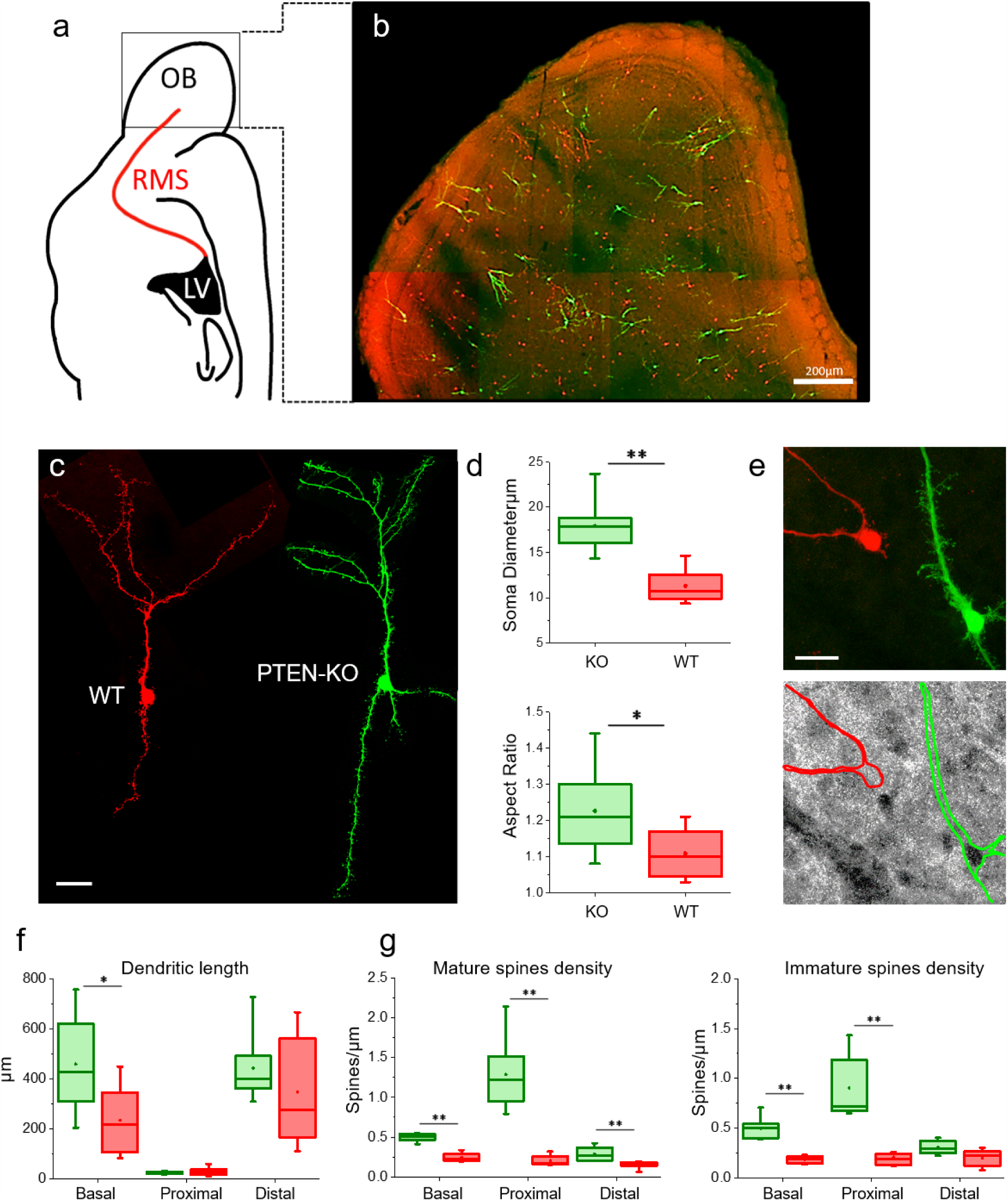
PTEN-KO mosaic of olfactory granule cells generated post-natally. **a**, Granule cell continuously differentiate during post natal life from progenitors lining the surface of the lateral ventricle (LV). New born neurons are electroporated at P3 and migrates following the rostral migratory stream (RMS) toward the olfactory bulb (OB). **b**, The co-transfection of Beatrix with a low titer of Cre plasmid causes the formation of a sparse mosaic of PTEN expression of GABAergic interneurons into the granule cell layer (imaging performed at P30). Scale bar 0.2 mm. **c**, Comparison between a WT and a PTEN-KO neuron in the olfactory bulb. Scale bar 50 μm. **d**, The loss of PTEN caused the hypertrophy of the cell body and altered its morphology. **e**, Immunohistochemistry against PTEN confirms the loss of the protein upon Beatrix recombination. Scale bar 20 μm **f**, The structure of the dendritic arborization is affected by the loss of PTEN as demonstrated by the length of the dendritic compartments. **g**, Spine density is altered both for mature spines (left) and for immature filopodia (right). Interestingly, this effect has a different spatial pattern than observed in pyramidal neurons (see Fig 3e). (n neurons: 9 WT, 9 KO; n spines: 1918 WT, 6,205 KO; Mann-Whitney statistical test).

**Supplementary Fig. 7.**
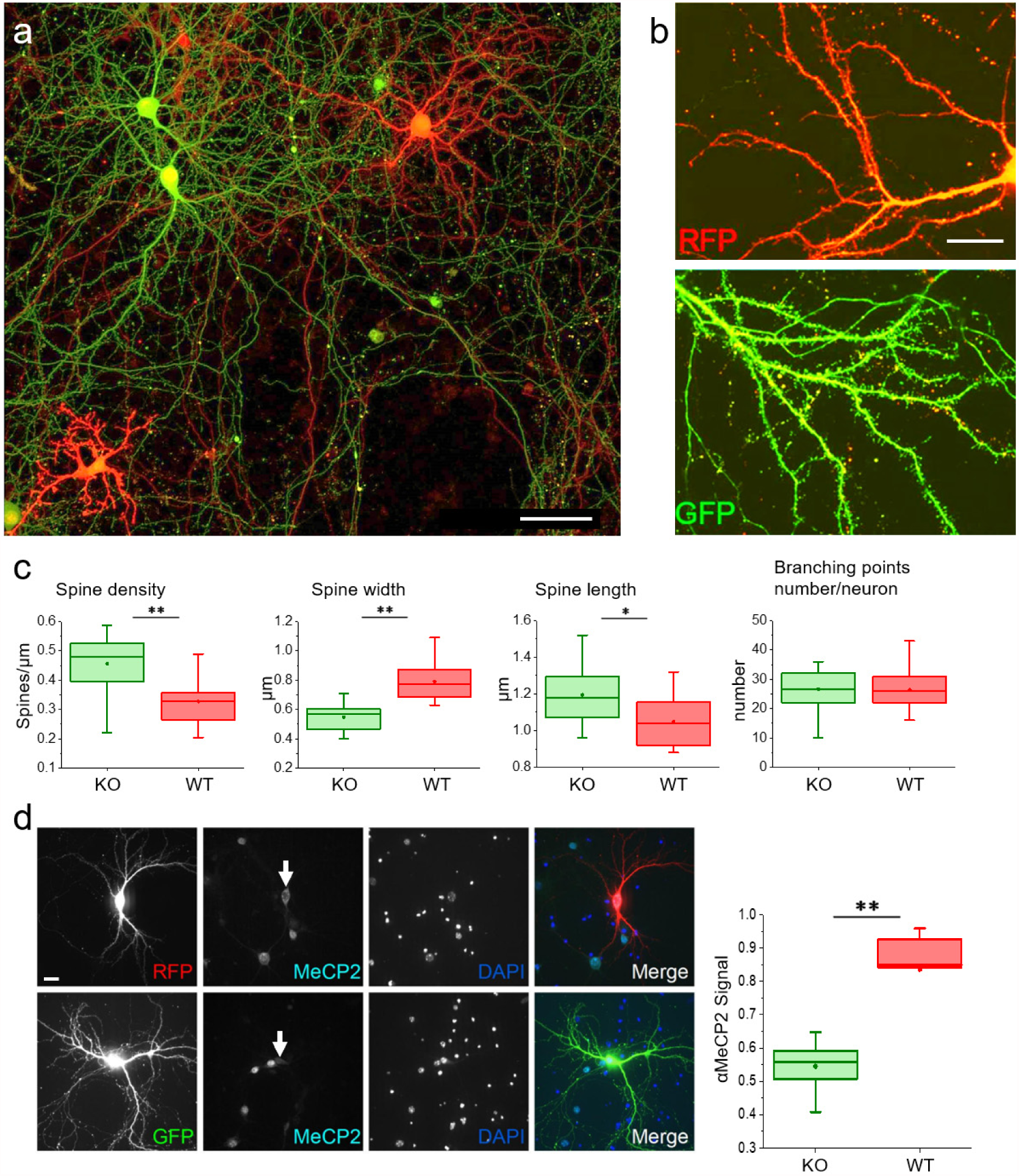
Cell-autonomous alterations of dendritic spines in a MeCP2 mosaics in neuronal cultures. **a**, Sparse MeCP2 KO mosaics at 12DIV in dissociated mouse cortical neuron cultures obtained from MeCP2^flox^ mice. Scale bar 50 μm **b**, High magnification details of dendrites from WT (top) and MeCP2-KO neurons (bottom). Scale bar 10 μm **c**, MeCP2 ablation in cultured cortical neurons leads to increased spine density with respect to the internal controls. This increase in spine number is characterized by morphological alterations, with spines presenting an average reduction of the width and a general increase in length. No difference has been observed between KO and WT neurons in the total number of branching points. (n neurons: 14 WT, 18 KO; Mann-Whitney statistical test?). **d**, MeCP2 immunostaining quantification (right) confirms the loss of MeCP2 in GFP^+^ cells, an its presence in RFP^+^ cells in the mosaics generated in cultured neurons. These data confirms the possibility to extend the use of Beatrix-induced mosaics in a different experimental paradigm, including a different genomic target. Scale bar 20 μm.

**Supplementary Fig. 8.**
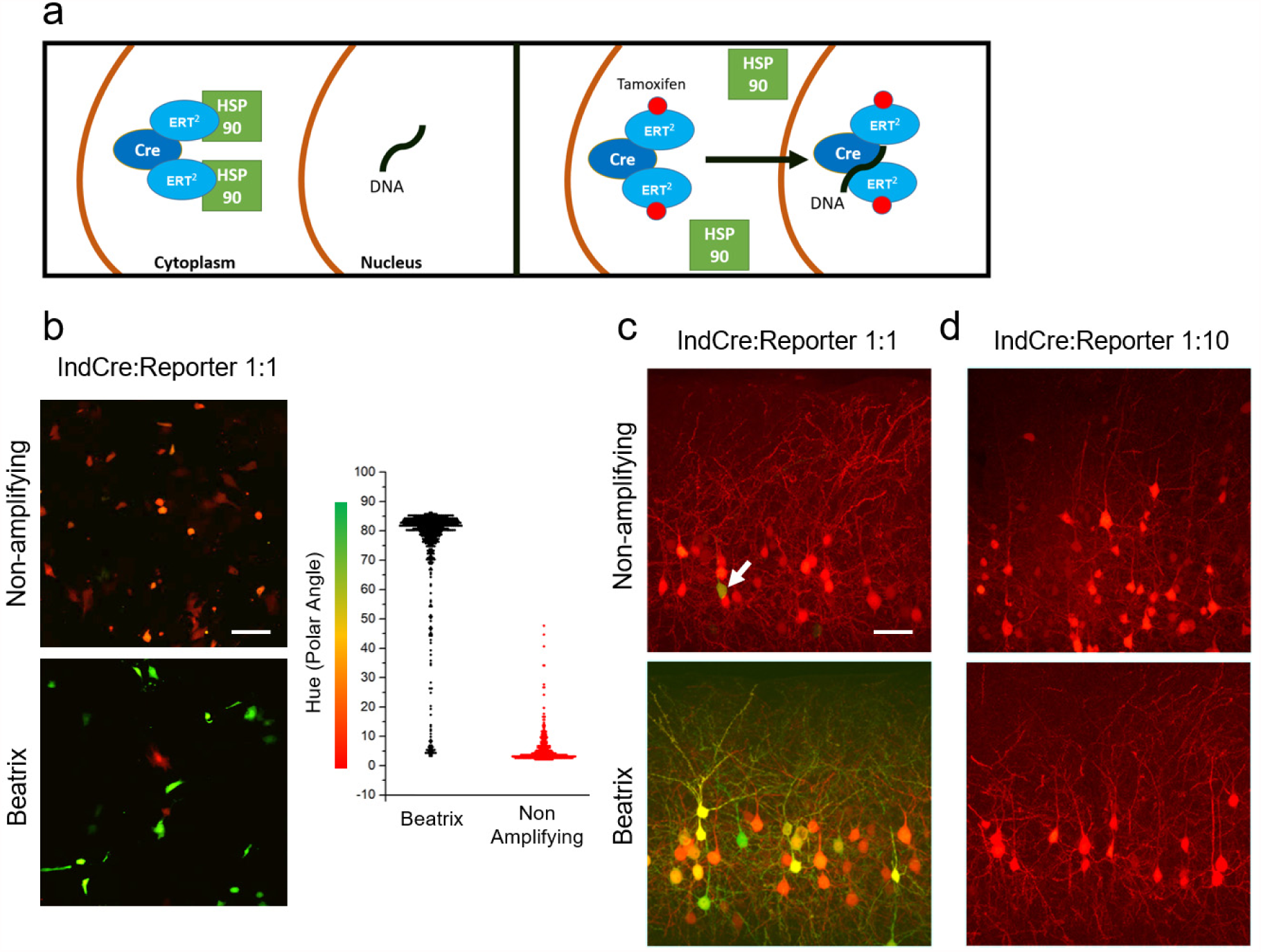
Inducible Cre leakage is unmasked by Beatrix high sensitivity. **a**, ERT2-Cre-ERT2 is an inducible Cre form with the lowest reported leakage levels (Matsuda and Cepko 2007). ERT2-Cre-ERT2 is kept outside the nucleus by the interaction with HSP90 that acts as a cytosolic anchor. In presence of tamoxifen, the anchor is released and ERT2-Cre-ERT2 is free to shuttle inside the nucleus leading to Cre-mediated DNA recombination. The sequestration of ERT2-Cre-ERT2 depends on the dissociation constant of the protein and on two independent reversible binding events with tamoxifen. Because of the self amplifying nature of Beatrix, this system has a very high sensitivity to low Cre-activity levels and this leads to leakage. **b**, The co-transfection of Beatrix with ERT2-Cre-ERT2 in 1:1 ratio revealed a large degree of leakage as shown in HEK 293T cells 4 days after transfection (bottom panel). In contrast, a non-amplifying Cre reporter is blind to leakage (top). The plot on the right demonstrates that most of the Beatrix-transfected cells are recombined, in stark contrast to cells transfected with the non-amplifying plasmid. **c**, A similar result has been obtained in vivo after *in utero* electroporation of ERT2-Cre-ERT2 together with Beatrix or with a non-amplifying reporter in a 1:1 ratio. The cortex transfected with the non-amplifying reported showed very limited leakage (white arrowhead, n=2 mice), while Beatrix show a large degree of leakage that accumulated in the time interval between electroporation and imaging (n=3 mice). Data acquired at P45. **d**, The leakage present in the Beatrix cortex can be effectively countered by lowering the concentration of the ERT2-Cre-ERT2 plasmid at the time of transfection to ratio 1:10 (n=4 mice; n=4 mice). Scale bar 50 μm.

**Supplementary Fig. 9.**
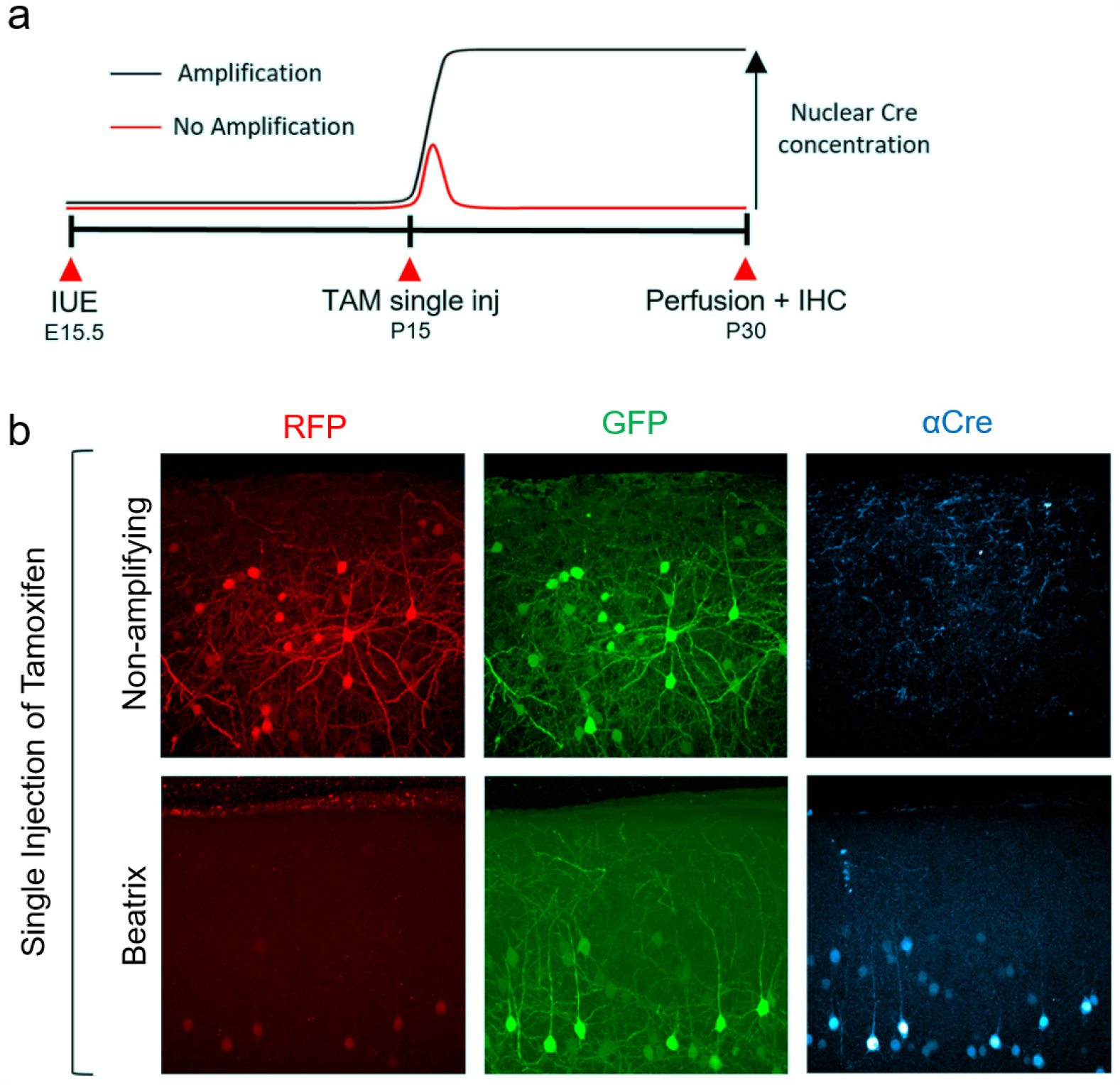
Beatrix amplifies and preserves the effects of a single pulse of tamoxifen *in vivo*. **a**, Timeline of the experiment. ERT2-Cre-ERT2 was electroporated *in utero* together with Beatrix or the non-amplifying reporter in a 1:10 ratio. At P15 we performed a single tamoxifen injection (100 μg/g) that usually does not lead to complete recombination (Hannah et al. 2018). **b**, A single tamoxifen injection at P15 led to a transient Cre activation in most cells, but, as witnessed by the co-expression of RFP, the reporter was not fully recombined thus proving an ambiguous reporting of the recombination state (n=3 mice). In presence of Beatrix, every cell respond to the transient Cre activation by triggering the self amplification, thus leading to complete recombination and to the loss of the red fluorescence (n=5 mice). Cells with a high level of expression show some residual red fluorescence localized in the nucleus because of the long halftime of dsRed2 protein in cells (Verkhusha et al. 2003). The stabilization of the transient Cre activation was confirmed by immunofluorescence staining of Cre protein, attesting the presence of the enzyme in cell nuclei 15 days after tamoxifen injection (bottom panel). On the other side, no Cre-positive cells were found in animals electroporated with the inducible Cre and the non-amplifying reporter (upper panel), indicating how the recombinase activity in this case is temporally confined to the 24-36h following the injection, thus leading to the massive presence of non-recombined or partially recombined cells.

## Materials and Methods

### Generation of the pCAG-Beatrix plasmid

We designed a custom plasmid (pUC-flex-GFP-P2A-Cre ex1) containing the eGFP cassette, the Lox sequences and Cre exon1. This plasmid was produced by the Genewiz gene synthesis facility (South Plainfield NJ, USA). The Cre exon2 was cloned upon amplification from a Cre ORF plasmid using BssHII and SacI sequences to obtain the pUC-flex-GFP-P2A-Cre ex1-ex2 plasmid. Then, we inserted the dsRed expression cassette into the AflII/XbaI sites, between the first LoxN and Lox2272 sites of pUC-flex-GFP-P2A-Cre ex1-ex2. This cloning step finally yield the pUC-dsRed-flex-GFP-P2A-Cre ex1-ex2 plasmid. The entire cassette was subsequently moved into an expression vector for *in utero* electroporation (Addgene #13777)^21^, downstream the CAG promoter sequence using EcoRI and NotI, to generate the final pCAG-Beatrix structures used in all experiments. For the generation of pCAG-BeatrixNE2 plasmid, the Cre exon2 in pCAG-Beatrix has been removed by substituting it with an empty DNA from pUC-flex-GFP-P2A-Cre ex1, using BsrGI and SacI restriction sites. All the DNA used for cloning and transfection was produced by using the QIAGEN Plasmid Plus Maxi Kit. Transformations have been performed in One Shot Stbl3 chemically competent *E. coli* cell line (Thermo Fisher Scientific), to avoid unwanted recombination events during plasmid preparation.

### Cell culture transfection and plasmid extraction

HEK293T and NIH3T3 cells were cultured in Dulbecco Modified Eagle’s Medium (DMEM) supplemented with 1mM Sodium Pyruvate, 2mM L-glutamine, 10U/mL–10μg/mL Penicillin-Streptomycin, 10% bovine foetal serum, 10 mM HEPES. All transfections were performed through electroporation on cell suspension. In brief, cells cultured on P60 Petri dishes (60mm diameter) at 75% confluency, were resuspended in 130 μl of electrolytic buffer, together with 10 μg of plasmidic DNA. Resuspended cells were electroporated using MicroPorator MP-100 (Digital Bio, 2 pulses at 1200 V with a duration of 20 ms).

### Animals

Timed pregnant CD1 mice (strain code 022; Charles River) were obtained from Charles River Laboratories, Inc. (Wilmington MA, USA) and maintained at Istituto di Neuroscienze (CNR, Pisa) animal house on a 12 h light cycle and fed ad libitum. Animal gestational ages were determined and confirmed during surgery. C57Bl/6J –PTEN^Flox^ animals (Jackson C;129S4-Pten^tm1Hwu^/J) were timed crossbred 16 days before the IUE experiment, performed at E.15. To follow the pregnancy status, females were weighed at different time points: 15, 7 and 2 days before the experiment. Both male and female subjects were used for mosaics generation and analysis. All animal care and experimental procedures were performed in accordance with Centro di Biomedicina Sperimentale-Area della Ricerca CNR and CNR-Istituto di Neuroscienze licensing. All procedures and experimental approaches were performed in strict accordance with the recommendations of the Italian Ministry of Health (Dlg. 26/14) and according to protocol 277/2015-R approved by the Ministry of Health on April 23, 2015 and protocol CBS-PT.0417 emitted on November 2017 via WEB..

### *In utero* electroporation

Tripolar *in utero* electroporation targeting pyramidal neurons of the visual cortex was performed as previously described ^13,22^. The day of mating (limited to 4 h in the morning) was defined as embryonic day zero (E0), and the day of birth was defined as postnatal day zero (P0). E15 timed-pregnant mice (gestation period ~21 days) were anesthetized with isoflurane (induction, 4%; surgery, 1%), and the uterine horns were exposed by laparotomy. The DNA (3–4 μg/μl) together with the dye Fast Green (0.3 mg/ml; Sigma, St Louis, MO, USA) was injected (~1 μl) through the uterine wall into the right lateral ventricle of each embryos by a 33-gauge standard point hypodermic needles (Sigma-Aldrich, Germany). After soaking the uterine horn with a pre-warmed phosphate-buffered saline (PBS) solution, the embryo’s head was carefully held between tweezers-type circular electrodes (5 mm diameter), while the third electrode (7×6×1 mm, gold-plated copper) was accurately positioned on the occipital lobe. Electroporation was accomplished with a BTX ECM 830 pulse generator (BTX Harvard Apparatus) and BTX 47 tweezertrodes. Six electrical pulses (amplitude, 30 V; duration, 50 ms; intervals, 1 s) were used for electroporation. The uterine horns were returned into the abdominal cavity, and embryos continued their normal development until delivery. Success rate for electroporation experiments varied depending on the mouse strain. For wild type C57Bl/6J and CD1 animals the survival rate after treatment was among 70% and 75%, whereas for C57Bl/6J –PTEN^Flox^ animals was about 60%. In all these cases the electroporation was effective in more than 90% of the surviving treated animals.

### Postnatal electroporation

pCAG-Beatrix plasmid electroporation targeting neuronal precursors of interneurons in the subventricular zone (SVZ) was performed in mice at postnatal day 2-4. Pups were anesthetized by placing them in a glass Petri dish pre-chilled at 4°C and then placed in wet ice. The DNA (4 μg/μl) was injected (2 μl) with a Hamilton syringe into the lateral ventricle. The site of injection was approximately equidistant from the lambdoid suture and the eye, 2 mm lateral to the sagittal suture and 2 mm deep from the scalp surface^23^. Electrodes were placed on the lateral walls of the pup head and ten electrical square pulses (amplitude, 100 V; duration, 50 ms; intervals, 450 ms) were delivered (NEPA 21 type 2 electroporator, NEPAGENE). Electroporated pups were warmed on a heating pad before returning into the homecage. The survival rate after treatment for C57Bl/6J – PTEN^Flox^ animals was about 80%. The electroporation was effective in about 80% of the surviving treated animals.

### Animals and surgical procedures

Experiments were performed on CD1, C57Bl/6J– PTEN^Flox^ mice (males and females) between postnatal day 28 and 32. Mice were anesthetized with an intraperitoneal injection of Urethane (20% w/V in physiological saline, 20 mg/Kg Urethane). During the experiment the animal respiration was aided providing O_2_ enriched air and body temperature monitored and held constant at 37 °C with a feedback-controlled heating blanket (Harvard Instruments). The animal head was shaved and 2.5% lidocaine gel applied to the scalp. Scissors were used to cut the flap of skin covering the skull of both hemispheres; the exposed bone was washed with saline and periosteum was gently removed with a pair of forceps to provide a better adhesion for glue and dental cement. The area expressing the highest fluorescence was identified with transcranial epifluorescence illumination. A custommade steal head post with a central imaging chamber was then glued with cyanoacrylate in a plane approximately parallel with the skull over the cortical region of interest and cemented in place with white dental cement (Paladur). The mouse was head-fixed and a craniotomy of 2– 3mm in diameter was drilled over the region of interest; care was taken to minimize heating of the cortex during surgery, dural tears, or bleeding, and to keep the cortex superfused with sterile ACSF (126mM NaCl, 3mM KCl, 1.2 mM KH_2_PO_4_, 1.3 mM MgSO_4_, 26 mM NaHCO_3_, 2.4 mM CaCl_2_, 15 mM Glucose, 1.2 mM HEPES in distilled H_2_O, pH 7.4).

### *In vivo* two-photon image acquisition

Fluorescence was imaged with a Prairie Ultima Multiphoton microscope (Prairie Technologies/Bruker) and a mode-locked Ti:sapphire laser (Chameleon Ultra II, Coherent) through a 20× Olympus XLUMPLFLN water immersion objective (numerical aperture 1.0). Before each imaging session we measured the power of the excitation laser at the optic bench and at the output of the objective lens at each wavelength. From these data we computed the power at the sample, which is not accessible once the mouse is placed under the objective, in order to maintain it under 20 mW. Fluorescent cells were imaged at variable depth in the cortex. PMT gain was kept constant at 667 V since previous calibrations showed that this voltage gives the best compromise between S/N ratio and dynamic range. All two-photon imaging experiments have been performed at the excitation wavelength of 950 nm. The emission filters band-pass used were 490-560 nm for the green channel and 584-680 nm for the red channel. For cultured cells experiments, multiple snapshots at a resolution of 512×512 or 1024×1024 pixels and zoom 1, were taken 48 h after cell transfection. *In vivo* imaging, was performed by acquiring a set of Z-stacks in the transfected regions. Dark frames were acquired after closing the laser shutter to measure the mean noise arising in the PMTs and the pedestal usually added by the electronics. Time lapse imaging of dendritic structures was restricted to the apical dendrites present in cortical layers 2/3 (50–150 mm below the cortical surface), and was conducted for at least 50–60 min at an interval of 5 minutes. Images were acquired with a water immersion lens (Olympus, 20X NA) at a resolution of 512×512 pixels at zoom 10, leading to a field of about 50 μm and a nominal linear resolution of about 0.1 μm/pixel. The stack step size was 0.75 mm. The acquisition of each frame was synchronized to the hearth beat and this procedure strongly reduced the mechanical artefact produced by the pulsing brain circulation and improved the alignment of the image stacks^24^. Analysis of the imaging data was carried out using ImageJ. The optical sections of each stack were aligned to compensate for mechanical movements by using the SIFT ImageJ plugin^25^.

### Cell culture and transfection of primary mouse cortical neurons

Cultured mouse cortical neurons were prepared from E16.5 embryos of WT or MeCP2^flox^ mice (MMRRC; B6; 129S4-Mecp2^tm1Jae^/Mmucd). Neurons were plated at 150-200 cells/mm² density on 12-well plates (Euroclone) coated with 0.01 mg/ml poly-L-Lysine (Sigma-Aldrich) and manteined in Neurobasal (ThermoFisher) supplemented with B27 (home made). At day-in-vitro 7 (DIV7), neurons were transfected using Lipofectamine 2000 (ThermoFisher). Imaging experiments were performed at DIV14.

### Immunocytochemistry

Cells were fixed in 4% paraformaldehyde, 4% sucrose in PBS [136.8 mM NaCl, 2.68 mM KCl, 10.1 mM Na_2_HPO_4_ and 1.76 mM KH_2_PO_4_, pH 7.4 (all Sigma-Aldrich)] at room temperature for 10 minutes. Primary antibodies were dissolved in homemade gelatin detergent buffer (GDB) [30 mM phosphate buffer, pH 7.4, 0.2% gelatin, 0.5% Triton X-100, 0.8 M NaCl (all Sigma-Aldrich)] and applied o/n at 4°C. Secondary antibodies conjugate with fluorophores (Jackson ImmunoResearch Laboratories) were also dissolved in GDB buffer and applied for 1h. DAPI staining (ThermoFisher) Coverslips were mounted using Mowiol mounting medium.

### Image acquisition and processing of cultured neurons

Confocal images were obtained using LSM 510 Meta confocal microscope (Carl Zeiss, gift from Fondazione Monzino) with Zeiss 63X or 25X objectives at a resolution of 1024×1024 pixels. Images represent averaged intensity Z-series projections of 2-7 individual images taken at depth intervals of around 0.45 μm. For dendritic spine analysis, morphometric measurements were performed using Fiji/ImageJ software (US National Institutes of Health). Individual dendrites were selected randomly, and their spines were traced manually. Maximum length and head width of each spine were measured and archived automatically.

### Immunohistochemistry

Animals were deeply anesthetized with urethane and perfused transcardially with 4% paraformaldehyde/PBS (4% PFA). Samples were post fixed overnight in 4% PFA. For immunofluorescence, brains were sectioned at 50 µm (100 µm for OB sections) thickness on a vibratome (Leica VT 1000S). Sections were processed as free-floating sections and stained with, GFP (Abcam, ab6673) RFP (Abcam, ab62341) Cre (Sigma-Aldrich, C7988) and PTEN (Cell Signaling, 9559) antibodies. Secondary immunostainings were performed using Alexa Fluor conjugates secondary antibodies (Abcam AF405, AF647).

### Confocal image acquisition of brain slices

Fixed tissue was imaged with the Zeiss LSM-800 Airyscan confocal microscope with 405/488/561/640 nm lasers according to the secondary antibody. Low-magnification images were acquired using a ×20 air objective (NA 0.5). Representative fields from thick slices (50 μm) were imaged by acquiring a set of Z-stacks (50 μm thick) for each experimental condition. The *Z*-stacks were collapsed with a maximum intensity projection to a 2D representation. Similarly, high-magnification images were acquired with a ×60 oil immersion objective (NA 1.3), and post processed to extract the details of the cellular morphology (dendrites and spines).

### Two-photon imaging analysis

Two-photon data were composed by 12bit images and have been analyzed with ImageJ (NIH). For cell culture, after background subtraction, images were analyzed using a custom-made ImageJ macro function to automatically identify regions of interest (ROIs) around single cells (see appendix). The macro computes the average fluorescence of each cell. *In vivo* Z-series were projected (maximum intensity) and corrected for dark background. Regions of Interest (ROIs) were manually drew around cellular somata: all pixels within each ROI were averaged to give the fluorescence value. Imaging data were not corrected for bleed through of emission in the two emission channels. Polar angle and radial distance were calculated from the colorimetric plots as previously described in paragraph 2.5.

Confocal acquisition were composed of 16bit images and were analyzed with ImageJ. Z-series for colocalization experiments, were performed through a frame by frame pixel intensity spatial correlation analysis with the Coloc2 ImageJ plugin function. For anti-colocalization experiments, the same procedure has been applied between RFP/GFP positive pixels and 0 value pixel from the AF647 Alexa Fluor conjugate immunostaining, obtained upon background subtraction. For display purposes, all images presented in the study received an identical non-linear stretching to improve visibility of the fainter details.

